# 3D osteocyte lacunar morphometry of human bone biopsies with high resolution microCT: from monoclonal gammopathy to newly diagnosed multiple myeloma

**DOI:** 10.1101/2023.07.12.548656

**Authors:** Inés Moreno-Jiménez, Sharen Heinig, Unai Heras, Daniela Simone Maichl, Susanne Strifler, Ellen Leich, Stéphane Blouin, Peter Fratzl, Nadja Fratzl-Zelman, Franziska Jundt, Amaia Cipitria

**Author notes:** These authors contributed equally to this work (Franziska Jundt, Amaia Cipitria).

## Abstract

Osteocytes are mechanosensitive, bone-embedded cells which are connected via dendrites in a lacuno-canalicular network and regulate bone resorption and formation balance. Alterations in osteocyte lacunar volume, shape and density have been identified in conditions of aging, osteoporosis and osteolytic bone metastasis, indicating patterns of impaired bone remodeling, osteolysis and disease progression. Osteolytic bone disease is a hallmark of the hematologic malignancy multiple myeloma (MM), in which monoclonal plasma cells in the bone marrow disrupt the bone homeostasis and induce excessive resorption at local and distant sites. Qualitative and quantitative changes in the 3D osteocyte lacunar morphometry have not yet been evaluated in MM, nor in the precursor conditions monoclonal gammopathy of undetermined significance (MGUS) and smoldering multiple myeloma (SMM). In this study, we characterized the osteocyte lacunar morphology in trabecular bone of the iliac crest at the ultrastructural level using high resolution microCT in human bone biopsy samples of three MGUS, two SMM and six newly diagnosed MM. In MGUS, SMM and MM we found a trend for lower lacunar density and a shift towards larger lacunae with disease progression (higher 50% cutoff of the lacunar volume cumulative distribution) in the small osteocyte lacunae 20-900 μm^3^ range compared to control samples. In the larger lacunae 900-3000 μm^3^ range, we detected significantly higher lacunar density and microporosity in the MM group compared to the MGUS/SMM group. Regarding the shape distribution, the MGUS/SMM group showed a trend for flatter, more elongated and anisotropic osteocyte lacunae compared to the control group. Altogether, our findings suggest that osteocytes in human MM bone disease undergo changes in their lacunae density, volume and shape, which could be an indicator for osteolysis and disease progression. Future studies are needed to understand whether alterations of the lacunae architecture affect the mechanoresponsiveness of osteocytes and ultimately bone adaptation and fracture resistance in MM and its precursors conditions.

## 1. Introduction

Osteocytes are bone-embedded cells, connected via dendrites in a lacuno-canalicular network (LCN). They act as local sentinels in response to external stimuli and regulate the balance between bone resorption/formation ^1^. In response to mechanical strain, osteocytes orchestrate the remodeling of the perilacunar bone matrix and thereby change shape and size of their lacunae ^2^. Lacunar volume, shape and density were influenced by changes of the surrounding environment, either mechanically or chemically, and by aging or disease. Changes in lacuna volume were observed after exposing mice to non-gravity ^3^ or after treatment with glucocorticoid slowing down bone metabolism ^4^, resulting in both cases in larger lacunae. In contrast, smaller lacunar size and lacunar density were observed in rodent models of aging ^5^ and osteoporosis, where the effect of different anti-resorptive drugs in intracortical microporosities was evaluated ^6^. In humans, a decline in lacunar density was observed with aging, while a significantly higher density was found in osteoporotic bone compared to controls ^7^. A recent study with human bone biopsy samples of healthy individuals across the life span revealed decreasing osteocyte lacunae density and porosity with aging, before the peak bone mass at 30 years, which then stabilized or even increased after this age ^8^. Furthermore, mineralized osteocyte lacunae (micropetrosis) were found increased with aging and osteoporosis, which was attributed to osteocyte apoptosis and improved with bisphosphonates treatment ^9^. Changes in lacunar shape, notably becoming smaller and more spherical with age, were linked to aging, resulting in diminished local bone tissue strains, impacting bone mechanosensitivity and fracture resistance ^2,10^. Noteworthy, abnormal morphological features were also detected in a large-scale assessment of adolescent idiopathic scoliosis, using high resolution microCT ^11^.

In cancer, osteocytes, as well as osteoblasts and osteoclasts, communicate with tumor cells and this generates a microenvironment that supports cancer cell growth and stimulates bone destruction ^12^. Interactions between breast and prostate cancer cells and osteocytes influenced cancer cell migration and invasion potential and contributed to the establishment and progression of bone metastasis ^13,14^. Also, breast cancer cells induced osteocyte apoptosis and altered osteocyte function ^15^. Recently, a study of osteotropic cancers in rodents revealed that the spatial distribution of cortical bone microporosity, osteocyte lacunae volume and the LCN characteristics were influenced by the presence of tumor cells in osteolytic lesions ^16^. Taken together, changes in the volume and shape of osteocyte lacunae are an indicator of osteolysis in bone metastasis of osteotropic cancers.

An example of an incurable malignancy of the bone marrow and bone is multiple myeloma (MM), in which plasma cells producing monoclonal immunoglobulins disrupt the bone homeostasis towards excessive bone resorption ^17^. MM cells alter the bone marrow environment by releasing signals that promote osteoclast activity, while inhibiting osteoblast function ^18,19^. There is evidence that MM cells also target osteocytes by enhancing apoptosis and expression of sclerostin, an inhibitor of bone formation ^20^. Even though MM cells directly interact with osteocytes and vice versa, the consequences of these interactions on the local bone microstructure are largely unknown ^20^.

In the clinical setting, two pre-symptomatic stages precede MM: (i) an initial phase named monoclonal gammopathy of underdetermined significance’ (MGUS), which can progress in (ii) smoldering multiple myeloma (SMM). Both MGUS and SMM patients do not present MM clinical symptoms (C.R.A.B. symptoms of MM: hypercalcemia, renal insufficiency, anemia and/or osteolytic bone lesions^21^). In patients with MM, low-dose whole body computed tomography (CT) scans are routinely used to detect osteolytic lesions and fractures ^22^. It remains, however, poorly understood whether bone microstructural changes precede the osteolytic lesions.

To identify associations between changes in the bone microarchitecture and fractures, x-ray absorptiometry measuring bone density or low-dose CT scans were used in two clinical studies ^23,24^. In one study, low-dose CT scans showed reduced radiodensity, trabecular thickening and vertebral fractures, demonstrating that focal skeletal changes were significantly associated with further skeletal damages ^23^. Some earlier studies had attempted to study bone trabecular thickness and separation in MM patients; however, the resolution limit of clinical CT scans was an obstacle for this objective ^25,26^. Our group used advanced imaging techniques (high resolution microCT, synchrotron phase contrast-enhanced microCT, confocal laser scanning microscopy) to evaluate the bone microstructure in an early MM mouse model ^27^. We showed irregular shaped osteocyte lacunae, altered osteocyte LCN characteristics and local alterations of the extracellular matrix (ECM) with active erosion sites ^27^. Following on from this preclinical study, here we investigate human bone biopsies of MGUS, SMM and newly diagnosed MM patients with high resolution microCT for ultrastructural characterization of osteocyte lacunar morphology, to identify alterations that can be linked to osteolysis and disease progression.

## 2. Materials and methods

### 2.1 Patient sample collection and clinical evaluation

The purpose of this study was to determine whether changes in osteocyte lacunae volume and shape distribution could be detected in mineralized bone biopsy samples from patients with MGUS, SMM and newly diagnosed MM. Patients with MGUS (n=3), SMM (n=2) and symptomatic MM (n=6) were diagnosed following the revised international staging system ^22^. Clinical data from eleven patients including reports of the low-dose CT scans are given in Tables 1-3. Values in brackets show the normal range for the parameters. The investigators who performed the bench studies were blind to the patient information. This study was approved by the Ethics Committee of the University of Würzburg, Germany (76/13 and 82/20-am). Patients where included with an informed consent for the bone biopsy, with a known history of MGUS, SMM, symptomatic MM, lymphoma, leukemia, myelodysplastic syndrome, or myeloproliferative disorders. Patients receiving a bone biopsy based on other disease like a bone marrow carcinosis were excluded from the study. As control bone biopsy samples, two different data sets were used (Table 4). One Jamshidi bone biopsy sample (2 mm diameter) was obtained from a patient of the University Hospital Würzburg with myelodysplastic syndrome and no diagnosis of MGUS, SMM or MM. Three cylindrical transiliac samples (5 mm diameter) were obtained from an earlier study ^8^ on autopsy samples at the Ludwig Boltzmann-Institute of Osteology Vienna. Osteocyte lacunae characteristics were re-evaluated in the current study. For all control and disease bone biopsy samples, the trabecular bone compartment was analyzed and the cortical bone was excluded. Trabecular bone is the first contact region with MM cells and comprises the larger fraction of bone in Jamshidi biopsies. Cortical bone was not present in some biopsies, as we received left over tissue after diagnosis. Only biopsy samples containing sufficient amount of trabecular bone, which maintained sample integrity without loss of bone structure (bone debris) caused by the extraction procedure, were considered.

**Table 1.** Clinical data from MGUS patients.

| Parameters |  |  |  |
| --- | --- | --- | --- |
| biopsy number | MGUS 1 | MGUS 2 | MGUS 3 |
| age (years) | 71 | 60 | 68 |
| gender | f | m | f |
| immunoglobulin subtype | IgG lambda | IgG kappa | IgG kappa |
| r-ISS | not applicable | not applicable | not applicable |
| cytogenetic risk | not applicable | not applicable | not applicable |
| M-protein [g/L] | not detectable | not detectable | 1.2 |
| heavy chain [mg/dL, normal range] | 1382<br>(610-1616)* | 881<br>(610-1616) | 507<br>(610-1616) |
| involved serum free light chain (sFLC) [mg/L, normal range] | 23.89<br>(5.71-26.30) | 10.75<br>(3.30-19.40) | 215<br>(3.30-19.40) |
| ratio (involved / uninvolved sFLC) | 0.86 | 1.02 | 5.44 |
| bone marrow infiltration of CD138<sup>+</sup> cells [%] | <10 | <10 | <5 |
| CT findings of osteolytic lesion | none | none | none |
| hemoglobin [g/dL] | 12.9<br>(12-16) | 14.4<br>(13.5-16.9) | 12.7<br>(12-16) |
| creatinine [mg/dL] | 0.63<br>(0-0.95) | 1.06<br>(0-1.17) | 6.69<br>(0-0.95) |
| cytogenetics** | no aberration | no aberration | no aberration |
| calcium [mmol/L] | 2.2<br>(2.0-2.7) | 2.5<br>(2.0-2.7) | 2.3<br>(2.0-2.7) |
| alkaline phosphatase [U/L] | 51<br>(35-105) | 72<br>(35-105) | not available |
| inorganic phosphate [mmol/L] | 1.09<br>(0.87-1.45) | 1<br>(0.87-1.45) | 1.85<br>(0.87-1.45) |
| bone anti-resorption treatment (duration) | Denosumab (four months) | no | no |

\* Values in brackets show the normal range for the parameters, respectively
\*\* Cytogenetic alterations by fluorescence in situ hybridization (FISH)

**Table 2.**
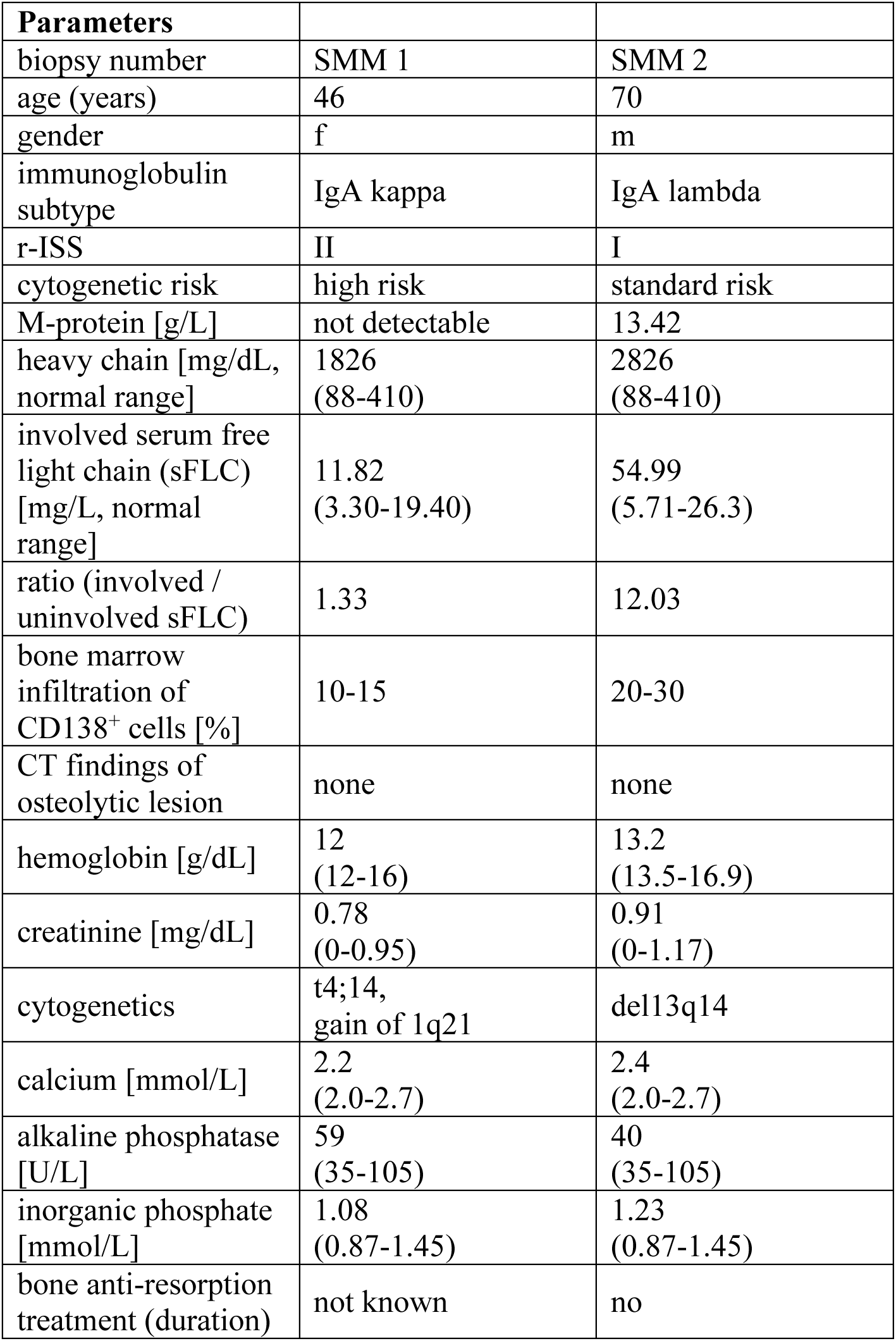
Clinical data from SMM patients.

**Table 3.** Clinical data from newly diagnosed MM patients.

| Parameters |  |  |  |
| --- | --- | --- | --- |
| biopsy number | MM 1 | MM 2 | MM 3 |
| age (years) | 58 | 57 | 72 |
| gender | m | m | f |
| immunoglobulin subtype | IgG kappa | kappa | IgG lambda |
| r-ISS | III | II | III |
| cytogenetic risk | high risk | standard risk | high risk |
| M-protein [g/L] | 35.3 | not detectable | 20.6 |
| heavy chain [mg/dL, normal range] | 4765<br>(610-1616) | not available | 3074<br>(610-1616) |
| involved serum free light chain (sFLC) [mg/L, normal range] | 71.12<br>(3.30-19.40) | 8260<br>(3.30-19.40) | 1376<br>(5.71-26.3) |
| ratio (involved / uninvolved sFLC) | 40.41 | 890.09 | 355.56 |
| bone marrow infiltration of CD138<sup>+</sup> cells [%] | 70 | 60-70 | 90 |
| CT findings of osteolytic lesion | osteolytic lesion with extraosseous soft tissue mass in spine (T12), lytic lesion in pelvis, rib series fractures, medullary lesions in femora and humeri, reduced bone mineral density | disseminated osteolysis in spine, one large lesion in T9 with increased fracture risk, old vertebral body collapse in T7 and fracture of the fourth left rib, multiple small osteolyses in scapulae, ribs, sternum reduced bone mineral density | no osteolytic lesions<br>medullary lesions in upper and lower limbs, spine, sternum; reduced bone mineral density |
| hemoglobin [g/dL] | 10.8<br>(13.5-16.9) | 10.7<br>(13.5-16.9) | 10.2<br>(12-16) |
| creatinine [mg/dL] | 0.83<br>(0-1.17) | 2.3<br>(0-1.17) | 0.72<br>(0-0.95) |
| cytogenetics | t14;16, gain of 1q21 | t11;14 | del 17p, 1q gain, del14q, del 4p, del 16q, del 5p |
| calcium [mmol/L] | 2.3<br>(2.0-2.7) | 2.9<br>(2.0-2.7) | 2.5<br>(2.0-2.7) |
| alkaline phosphatase [U/L] | 97<br>(35-105) | 80<br>(35-105) | 49<br>(35-105) |
| inorganic phosphate [mmol/L] | 1.25<br>(0.87-1.45) | 1.33<br>(0.87-1.45) | 0.99<br>(0.87-1.45) |
| bone anti-resorption treatment (duration) | bisphosphonate (zoledronic acid) | no | no |
|  | for one month |  |  |

**Table 3 (cont). Clinical data from newly diagnosed MM patients**
| Parameters |  |  |  |
| --- | --- | --- | --- |
| biopsy number | MM 4 | MM 5 | MM 6 |
| age (years) | 60 | 68 | 64 |
| gender | m | f | f |
| immunoglobulin subtype | IgA lambda | IgG kappa | IgG kappa |
| r-ISS | II | II | II |
| cytogenetic risk | standard risk | standard risk | standard risk |
| M-protein [g/L] | 32.6 | 4.5 | 7.4 |
| heavy chain (mg/dL, normal range) | 4000<br>(88-410) | 1180<br>(610-1616) | 1625<br>(610-1616) |
| involved serum free light chain (sFLC) [mg/L, normal range] | 33.8<br>(5.71-26.3) | 16.59<br>(3.30-19.40) | 309.8<br>(3.30-19.40) |
| ratio (involved / uninvolved sFLC) | 4.26 | 1.49 | 8.41 |
| bone marrow infiltration of CD138<sup>+</sup> cells [%] | 30 | 50 | 60 |
| CT findings of osteolytic lesion | osteolytic lesion in pelvis; mild reduction in bone mineral density | disseminated osteolysis in skull, spine, fracture series in spine (T7, 8, 11; L1, 2, 4), reduced bone mineral density in spine | none |
| hemoglobin [g/dL] | 14<br>(13.5-16.9) | 10.1<br>(12-16) | 12.8<br>(12-16) |
| creatinine [mg/dL] | 0.96<br>(0-1.17) | 1.02<br>(0-0.95) | 0.91<br>(0-0.95) |
| cytogenetics | del 1p32, del13q14, gain of 1q21 | trisomy 11 | gain of 1q21 |
| calcium [mmol/L] | 2.4<br>(2.0-2.7) | 2.5<br>(2.0-2.7) | 2.4<br>(2.0-2.7) |
| alkaline phosphatase [U/L] | 41<br>(35-105) | 95<br>(35-105) | 92<br>(35-105) |
| inorganic phosphate [mmol/L] | 1.19<br>(0.87-1.45) | 0.94<br>(0.87-1.45) | 0.74<br>(0.87-1.45) |
| bone anti-resorption treatment (duration) | no | no | no |

**Table 4.** Control samples: Jamshidi bone biopsy and samples from autopsies.

| Parameters |  |  |  |  |
| --- | --- | --- | --- | --- |
| biopsy number | control 1 | control 2 | control 3 | control 4 |
| type of biopsy | jamshidi | autopsy* | autopsy* | autopsy* |
| age (years) | 69 | 64 | 57 | 63 |
| gender | f | f | f | f |
| previous disease | myelodysplastic syndrome | not known | not known | not known |

\* Subset of an earlier cohort presented previously <sup>8</sup> and re-evaluated here

The patient clinical data (Tables 1-3) include: biopsy number, disease stage (MGUS, SMM or newly diagnosed MM), age, gender, immunoglobulin subtypes (heavy and light chains) secreted by malignant plasma cells, revised International Staging System (r-ISS), cytogenetic risk, M-protein secreted by plasma cells at time of diagnosis (g/L), heavy chain (mg/dL), involved serum free light chain (sFLC) (mg/L), ratio (involved/uninvolved sFLC), bone marrow infiltration (%) based on immunohistochemistry Syndecan-1 (CD138) with >60% corresponding to newly diagnosed MM, computed tomography findings of osteolytic lesions, hemoglobin (g/dL), creatinine (mg/dL) and cytogenetic alterations by fluorescence in situ hybridization (FISH), which allows for detection of genetic changes which are relevant for the ISS stage. In addition, parameters describing the bone metabolism include: calcium (mmol/L), alkaline phosphatase (U/L), phosphate (mmol/L) and finally, bone anti-resorption treatment ongoing at the time of biopsy.

### 2.2 Immunohistochemical detection of CD138 on decalcified bone paraffin sections

Paraffin embedded human specimens were retrospectively collected from formalin-fixed paraffin-embedded tissue of anonymized samples, archived at the Institute of Pathology, University of Würzburg. Shortly, Jamshidi bone biopsy samples that were derived from the clinical department of the University of Würzburg were fixed for 24 h in formalin, decalcified for 24 h at 37°C using an EDTA-based approach, dehydrated with alcohol and xylene and then embedded in paraffin.

For the immunohistochemical detection of CD138/Syndecan-1 expression, sections were deparaffinized. The tissue was hybridized with an antibody against CD138/Syndecan-1 (clone MI15, mouse, #M7228, Agilent, Santa Clara, CA, USA), following standard procedures by automated processing using the Tecan Evo Freedom stainer and the Dako Advance system according to the manufacturer’s protocol.

### 2.3 Microcomputed tomography (microCT): image acquisition

Specimens were scanned using an EasyTom micro/nano tomograph (RX solutions, France). Low resolution scanning parameters were: 107 kV, 93 µA, 7 µm voxel size, frame rate of 3, average frame of 5. High resolution scanning parameters were: 60 kV, 124 µA, 1-2 µm voxel size, frame rate of 1, average frame of 5. Reconstruction of the 1120 projections/scan were performed using RX Solutions X-Act software.

### 2.4 Low resolution microCT: image analysis

The overall bone biopsy morphology with particular focus on trabecular bone thickness was analyzed with Dragonfly software, Version 2022.1 for [Windows]. Comet Technologies Canada Inc., Montreal, Canada; software available at https://www.theobjects.com/dragonfly. Low resolution microCT stack images (voxel size of 7 μm) were segmented using Otsu thresholding (noise removal of these microCT images and objects was not necessary). A mesh was then created from each individual region of interest (ROI) of the trabecular bone. The trabecular bone strut thickness was automatically generated and measured from each mesh by Dragonfly software. For this, we calculated the median, first and third quartile, minimum and maximum values of about 3-5.5 10^6^ local vertices created between boundary points of each overall mesh. The software calculated these vertices as diameters of hypothetical spheres that fit within each boundary points of the mesh. The trabecular bone strut thickness distribution is depicted with a color code for each biopsy and as a box plot.

### 2.5 High resolution microCT: image analysis

The VOI within the high resolution scans were selected in such a way so that the bone volume to be analyzed was maximized. The empty cavities within mineralized tissue corresponding to osteocyte lacunae were segmented. Based on previous studies with healthy trabecular bone ^8,28–30^, cavities smaller than 20 μm^3^ and larger than 3000 μm^3^ were considered cracks or artifacts and excluded from the analysis.

#### 2.5.1 Segmentation of osteocyte lacunae

The stack of high resolution microCT images (1-2 μm voxel size) was segmented using Otsu thresholding in Dragonfly. For these images, noise removal was performed using 3D smoothing (kernel of 3) as morphological operation. Next, the binary mask was used to extract the pores. Osteocyte lacunae were filled in a copy of the binary mask using a filling morphological operation. Next, the multi-ROI conversion tool of Dragonfly was used to select each lacuna individually. This tool allowed to define each osteocyte lacuna as an individual ROI, hence enabling the calculation of each lacunae volume and shape. The generated binary images were then used for the quantitative analysis of lacunae volume and shape with Python software (version 3.5). The script used was written in a Jupyter notebook with Python software (version 3.7). External libraries not contained in the Anaconda environment were installed from their repository. Micropores smaller than 20 μm^3^ and larger than 3000 μm^3^ were considered cracks or artifacts and excluded from the analysis. For the purpose of this study, two separate lacunar volume ranges were investigated: small lacunae 20-900 µm^3^ (osteocyte lacunar microporosity characteristic of healthy bone ^28,31^), and larger lacunae 900-3000 µm^3^ (microporosity potentially associated to bone disease).

#### 2.5.2 Osteocyte lacunar volume analysis

For the osteocyte lacunar volume analysis, each segmented lacunar volume was calculated as the total number of voxels times the volume of a single voxel. Based on this, we could define the number of lacunae (Lc.N) of a given lacunar volume (Lc.V). We could also define the total number of osteocyte lacunae (Lc.TN), total lacunar volume (Lc.TV) and trabecular bone total volume (Tb.TV). This nomenclature and units are summarized in Table 5.

**Table 5.** Nomenclature for osteocyte lacunar volume analysis.

| Nomenclature | Abbreviation | Unit |
| --- | --- | --- |
| Number of osteocyte lacunae | Lc.N | - |
| Total number osteocyte lacunae | Lc.TN | - |
| Lacunar volume | Lc.V | $\mu\text{m}^3$ |
| Total lacunar volume | Lc.TV | $\mu\text{m}^3$ |
| Trabecular bone total volume | Tb.TV | $\mu\text{m}^3$ |

#### 2.5.3 Normalized lacunar volume frequency distribution and lacunar density

To plot the lacunar volume frequency distribution, all lacunae within one sample were stratified according to lacunar volume intervals. For a discrete lacunar volume interval of 20 μm^3^, we quantified the number of osteocyte lacunae (Lc.N.) for that given size. For a given lacunar volume (Lc.V), we quantified the lacunar number (Lc.N), normalized to the trabecular bone total volume (Tb.TV) and excluding the total lacunar volume (Tb.TV- Lc.TV), to account for size differences between samples, since larger samples have more lacunae than smaller ones, as well as for potential differences in the trabecular bone microporosity. The data is presented as the frequency in a histogram with a bin size of 20 μm^3^. Furthermore, the lacunar density was calculated as the total number of osteocyte lacunae (Lc.TN) normalized to the trabecular bone total volume and excluding lacunar total volume (Tb.TV-Lc.TV), in other words, the summation of the osteocyte lacunar volume frequency distribution. This was done for both ranges separately, small lacunae 20-900 µm^3^ and larger lacunae 900-3000 µm^3^.

#### 2.5.4 Lacunar volume cumulative distribution

The cumulative distribution is calculated by sorting all osteocyte lacunae by volume, from smallest to largest, and increasing the count of the fraction 1/N at each point, with N = length of array. The cumulative lacunae volume distribution was plotted with respect to the lacunae volume (Lc.V), with the x axis showing a given lacunae volume and the y axis showing the cumulative volume. The 50% cutoff illustrating the specific volume below which half of all osteocyte lacunae are contained, was marked with a dashed line.

#### 2.5.5 Osteocyte lacunar microporosity analysis

The microporosity was calculated as the lacunar total volume (Lc.TV) divided by trabecular bone total volume (Tb.TV) for each group. This was done for both ranges separately, small lacunae 20-900 µm^3^ and larger lacunae 900-3000 µm^3^.

#### 2.5.6 Osteocyte lacunar morphological analysis

To perform the lacunar morphological analysis, the resulting shape of a lacuna and model representation consists of a 3×3 matrix of the positions of the voxels in a lacuna. For a diagonal symmetric matrix defining an ellipsoid, this is converted into the eigenvectors of the matrix, which correspond to the major and minor axes of the ellipsoid. From the eigenvectors, the corresponding eigenvalues (EV) were derived (EV1 > EV2 > EV3), where EV1 is the largest eigenvalue and EV3 the smallest ^32,33^. Three parameters were assessed to examine lacunae morphology, based on the eigenvalues of each individual lacuna: the lacuna elongation, flatness and anisotropy ^34^. This nomenclature and units are summarized in Table 6. In short, lacuna elongation (EV2/EV1, ratio of medium to largest eigenvalue) ranged from lower values for elongated and 1 for spherical; lacuna flatness (EV3/EV2, ratio of smallest to medium eigenvalue) ranged from 0 for flat to 1 for spherical; and lacuna anisotropy (1- EV3/EV1, one minus the ratio of smallest to largest eigenvalue) ranged from 0 for isotropic/spherical to 1 for highest anisotropy.

**Table 6:** Nomenclature for osteocyte lacunar analysis.

| Parameter | Definition | Unit | Formula |
| --- | --- | --- | --- |
| Normalized lacunar volume frequency distribution | Number of lacunae volume divided by the trabecular bone total volume minus the lacuna total volume | $\#/\mu\text{m}^3$ | $\frac{\sum Lc.N}{Tb.TV - Lc.TV}$ |
| Lacunar volume cumulative distribution | Sorted sum of volume divided by total volume | - |  |
| Microporosity | Total lacunar volume with respect to the trabecular bone total volume | - | $\frac{Lc.TV}{Tb.TV}$ |
| Lacuna elongation | 0 for elongated<br>1 for spherical | - | $\frac{EV2}{EV1}$ |
| Lacuna flatness | 0 for flat<br>1 for spherical | - | $\frac{EV3}{EV2}$ |
| Lacuna anisotropy | 0 for isotropic, spherical<br>1 for highly anisotropic | - | $1 - \frac{EV3}{EV1}$ |

### 2.6 Statistical analysis

A total of n=15 bone biopsy samples were analyzed, including MGUS n=3, SMM n=2, newly diagnosed MM n=6 and control biopsy samples n=4 (Jamshidi n=1 and from autopsies n=3). Additional bone biopsy samples (n>15) were collected but excluded from the analysis due to lack of sample integrity (e.g. loss of bone structure (bone debris) during the extraction procedure) or insufficient amount of trabecular bone. To analyze the differences in bone microporosity and lacunar morphological analysis, a one-way analysis of variance (ANOVA) with Tukey’s multiple comparisons test (*p<0.05) was used (GraphPad Prism 8.0 software).

## 3. Results

### 3.1 Clinical data and patient history

Clinical data from patients from the University Hospital Würzburg and parameters describing bone metabolism are compiled in Tables 1-3. The cohort included patient-derived bone biopsy samples with the precursor condition MGUS (Table 1), SMM (Table 2), and newly diagnosed MM (Table 3). MGUS is diagnosed when monoclonal immunoglobulins (< 3 g/dL) are detected in the serum of patients and less than 10% abnormal plasma cells are found in the bone marrow. When MGUS continuously progresses, the condition is named SMM, when between 10-60% plasma cells are detected and monoclonal immunoglobulin levels are higher than 3 g/dL. SMM differs from symptomatic MM, a stage in which C.R.A.B. symptoms of hypercalcemia, renal insufficiency, anemia and osteolytic bone lesions occur ^35^. The revised International Staging System for Multiple myeloma (r-ISS) was used to stratify patients with respect to their survival (Tables 1-3) ^36^. Control samples obtained from the University Hospital Würzburg and from the Ludwig Boltzmann-Institute of Osteology Vienna are compiled in Table 4.

Patients with MGUS showed no detectable osteolytic bone lesions by low-dose CT scans (Table 1). One patient with MGUS received denosumab for four months (MGUS1, Table 1). In this case, we cannot exclude that the anti-resorptive treatment interfered with the bone ultrastructure. Similarly, the two patients classified as SMM exhibited no signs of osteolytic lesions by low-dose CT scans at the time of biopsy (Table 2). Disease staging for SMM was determined based on the percent of bone marrow infiltration of CD138^+^ plasma cells, above 10%. Patients classified as MM had a bone marrow infiltration of CD138^+^ plasma cells between 60-90% (4 out of 6), a ratio of the involved to the uninvolved free light chain in the serum (sFLC) of >100 (2 out of 6), and/or local or disseminated lytic lesions, as well as reduced bone mineral density (4 out of 6 patients) based on CT findings (Table 3). All bone turnover markers (phosphate, calcium, alkaline phosphatase) were within normal ranges (Tables 1-3). Anti-resorptive therapy with the bisphosphonate zoledronic acid had been administered to one MM patient for one month before the biopsy was taken (MM1, Table 3). We assume that the single dose of the bisphosphonate did not affect bone ultrastructure in such short timeframe.

### 3.2 Immunohistochemical (IHC) and microcomputed tomography (microCT) overview analysis

As part of the standard clinical practice, and to detect the clonal bone marrow plasma cells and quantify the percent of bone marrow infiltration of CD138^+^ cells, bone biopsies were processed for immunohistochemical (IHC) analysis for CD138 staining. Fig. 1a-c shows representative samples of MGUS, SMM and newly diagnosed MM patients with <10% for MGUS and >10% for SMM and MM patients. Biopsies collected from patients were selected after applying exclusion criteria (see section 4.6 for details). 3D reconstructions of representative images are shown in Fig. 1d-f. Quantification of trabecular thickness distribution is shown in Fig. 1g-j and Supplementary Figure S1. No clear differences between MGUS, SMM and newly diagnosed MM were observed in the macroscopic trabecular bone architecture, which motivated the further analysis of bone ultrastructure with high resolution microCT.

**Figure 1.**
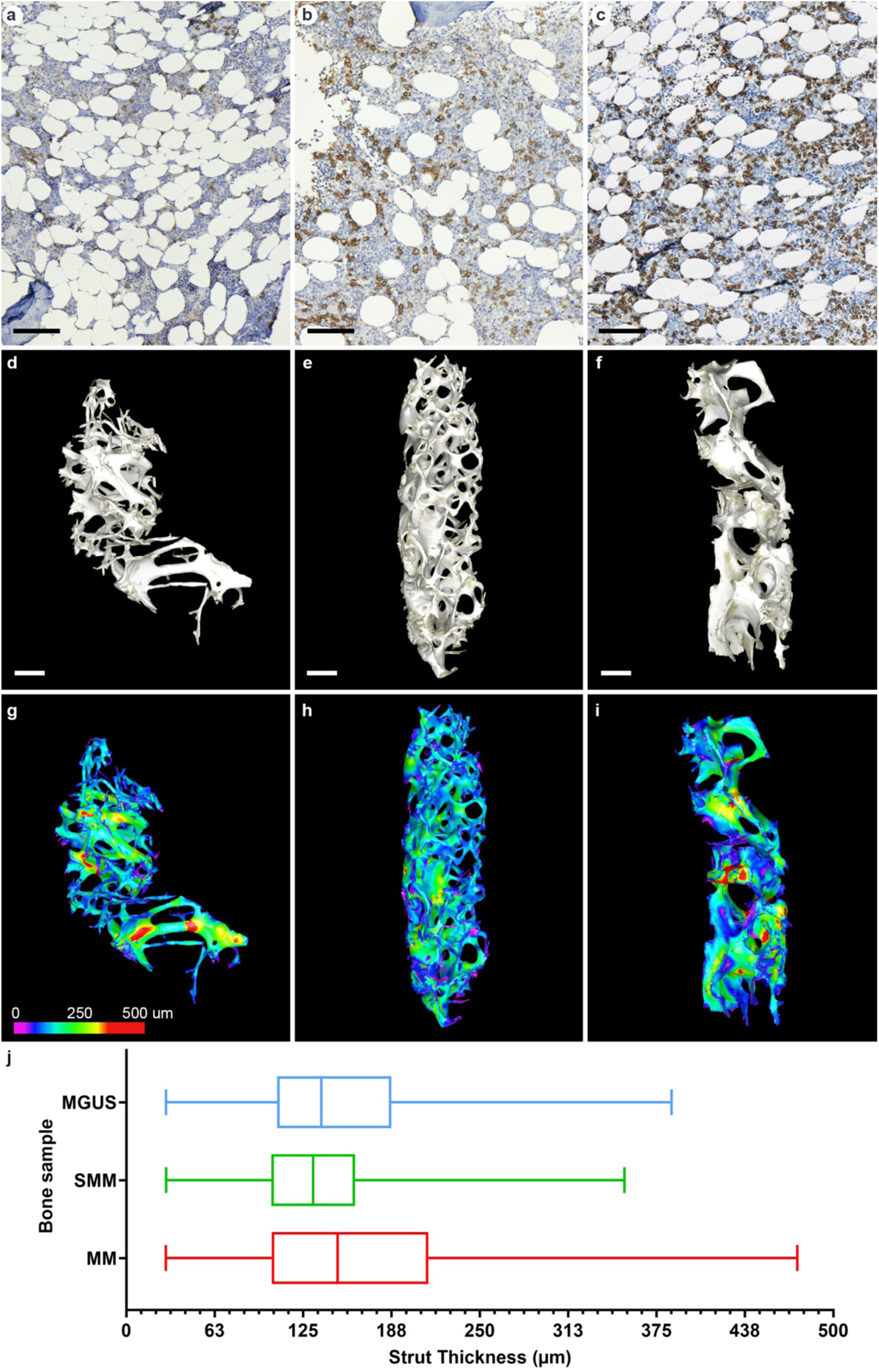
Immunohistochemical (IHC) CD138 staining and low resolution microCT overview of the bone biopsies from patients with: (a, d, g) monoclonal gammopathy of undetermined significance (MGUS), (b, e, h) smoldering myeloma (SMM), and (c, f, i) newly diagnosed multiple myeloma (MM). (a-c) Representative light microscopy images of the bone marrow compartment from the biopsy samples: CD138 positive staining in brown, cell nuclei counterstain in blue. MGUS bone biopsy #MGUS 2 with <10% bone marrow infiltration, SMM bone biopsy #SMM 1 with 10-15% bone marrow infiltration, MM bone biopsy #MM 1 with 70% bone marrow infiltration. (d-f) 3D reconstructions of low resolution (7 μm voxel) microCT scans of representative bone biopsies. (g-i) Quantitative analysis of trabecular bone strut thickness distribution and color code visualization with Dragonfly software. (j) Distribution of trabecular thickness shown with median, first and third quartile. Scale bars equal to 100 μm (a-c), 1 mm (d-f).

### 3.3 Bone ultrastructure and segmentation of osteocyte lacunar microporosity

We next investigated the bone ultrastructure with a focus on the osteocyte lacunar microporosity. High resolution microCT scans (voxel size 1-2 μm) were performed on smaller volumes of interest (VOI) (Supplementary Figure S2). Figure 2 illustrates an example of VOI of representative samples and the segmented osteocyte lacunar microporosity for MGUS, SMM and newly diagnosed MM, and Supplementary Figure S3 for a control sample. This segmented osteocyte lacunar microporosity was then quantitatively analyzed for lacunar volume distribution (Fig. 3-4), lacunar density and microporosity (Fig. 5), and lacunar morphological analysis (Fig. 6).

**Figure 2.**
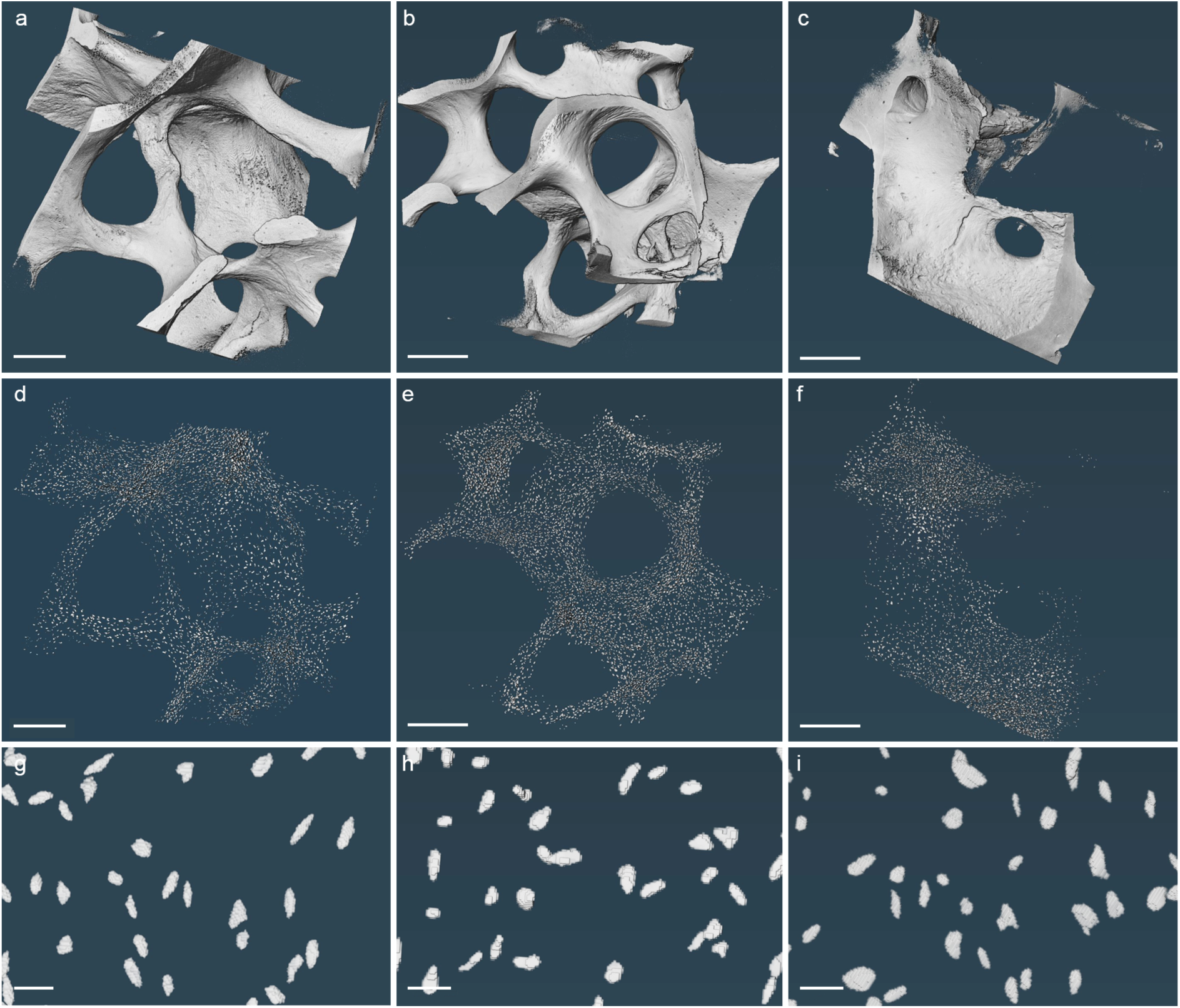
Bone biopsy macroscopic structure and osteocyte lacunar microporosity visualized with high resolution microCT in MGUS, SMM and MM samples. (a-c) 3D reconstructions of local scans (high resolution microCT; voxel size 1-2 μm) and (d-i) the corresponding segmented osteocyte lacunar microporosity of representative samples: (a, d, g) MGUS bone biopsy #MGUS 2, (b, e, h) SMM bone biopsy #SMM 1, (c, f, i) MM bone biopsy #MM 1. Scale bars equal to 250 μm (a, d), 350 μm (b, e), 400 μm (c, f) and 20 μm (g- i).

**Figure 3.**
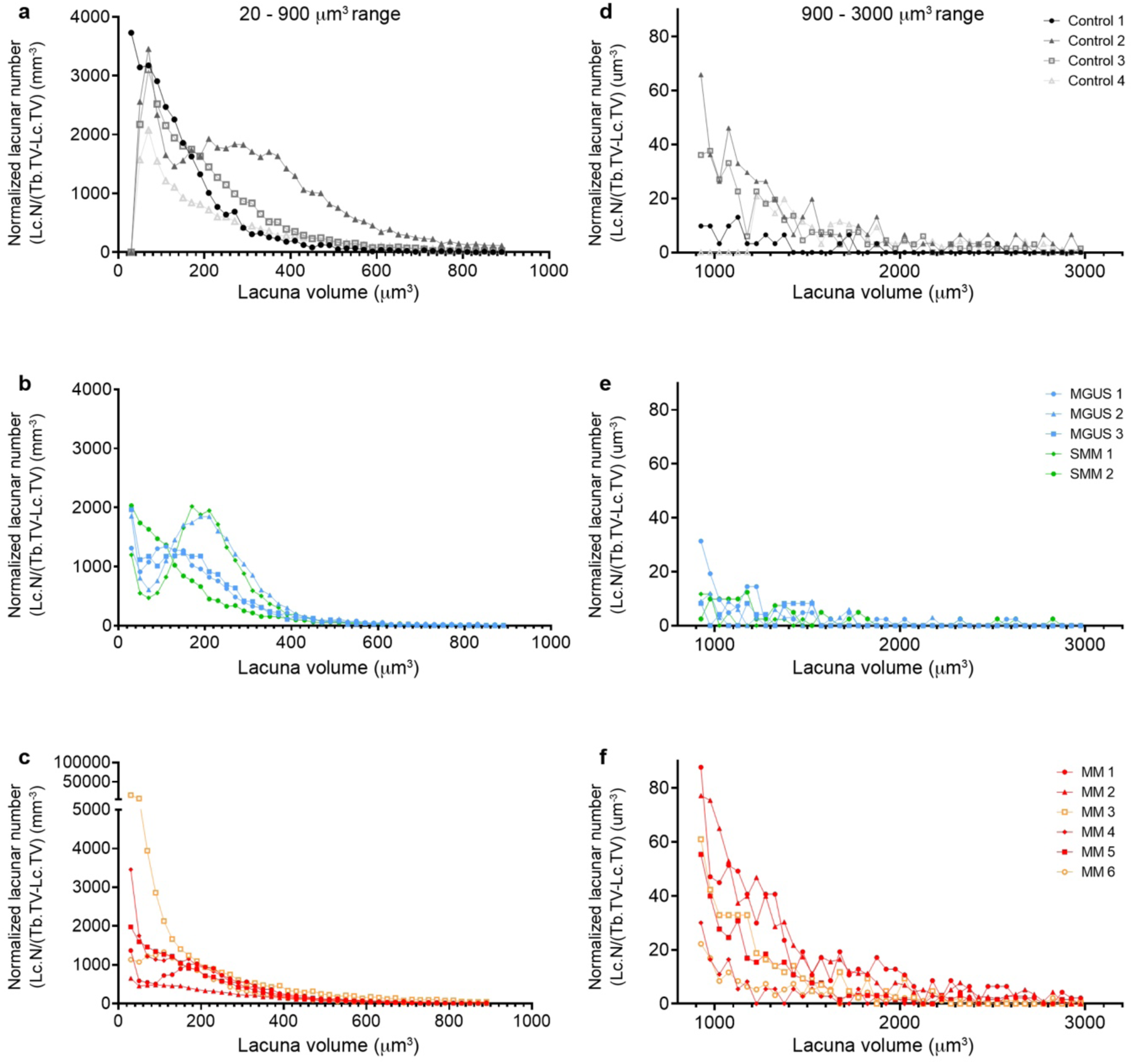
Lacunar volume frequency distribution of osteocytes in control, MGUS, SMM and MM samples. Frequency distribution normalized to trabecular bone total volume for two separate lacunar volume ranges: 20-900 µm^3^ and 900-3000 µm^3^. For each lacunae volume interval (x-axis), number of osteocyte lacunae (Lc.N) normalized by the trabecular bone total volume excluding lacunar total volume (Tb.TV-Lc-TV) (y-axis). Frequency distribution for each group: (a, d) control, (b, e) MGUS and SMM, (c, f) MM. Sample from the Jamshidi biopsy in black, all other control samples in grey. MGUS in blue, SMM in green, MM samples with bone lytic lesion in red, MM samples without lytic lesions in orange. Please note the different scale in the y-axis of subplot c.

**Figure 4.**
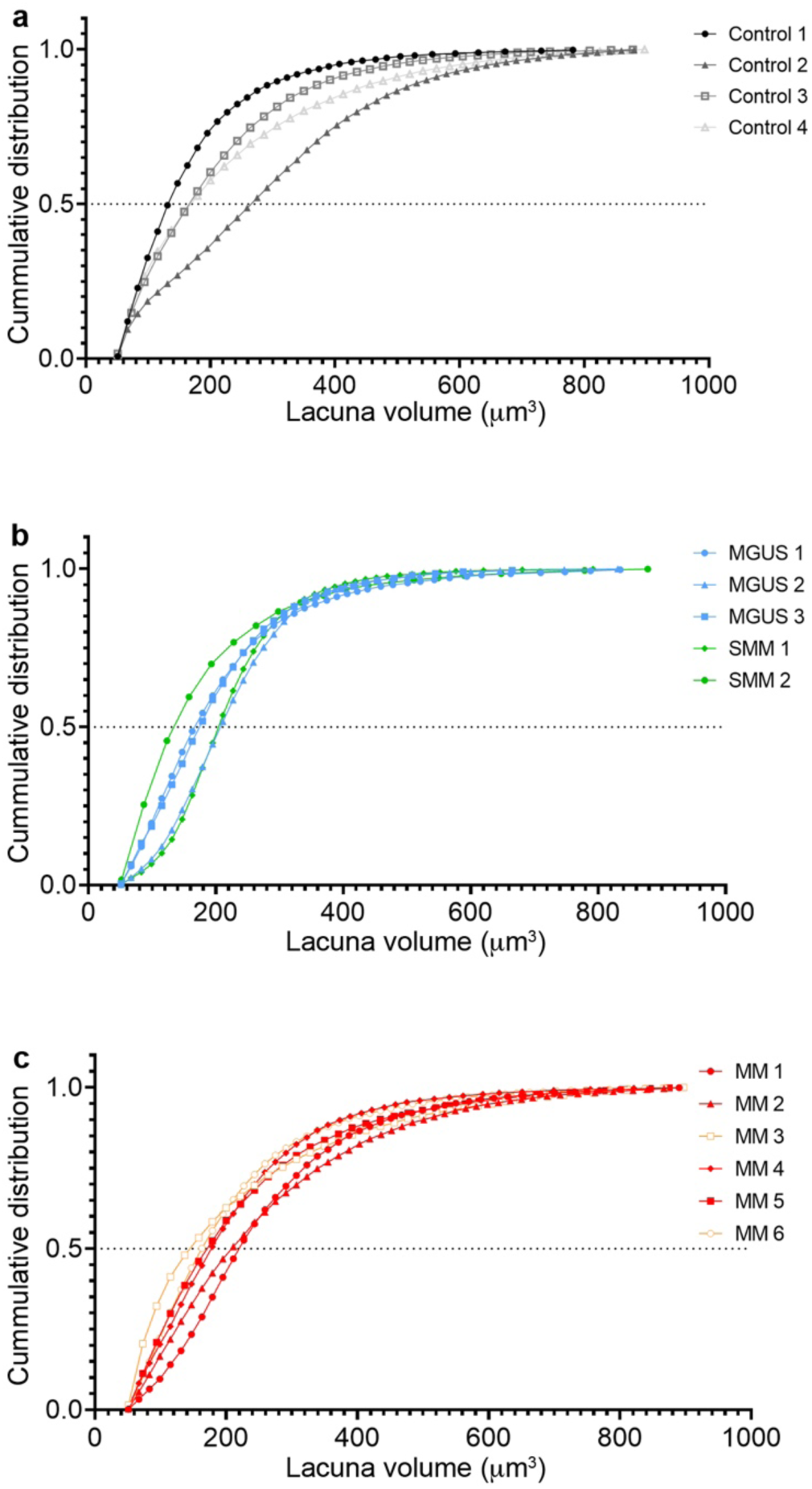
Lacunar volume cumulative distribution for the small lacunae 20-900 µm^3^ range of osteocytes in control, MGUS, SMM and MM samples. Cumulative distribution (from 0 to 1) of lacunae (y-axis) for each volume in µm^3^ (x-axis). Cumulative distribution for each group: (a) control, (b) MGUS and SMM, (c) MM. Dashed line indicates the 50% cutoff volume below which are half of all osteocyte lacunae of one sample. Sample from the Jamshidi biopsy in black, all other control samples in grey. MGUS in blue, SMM in green, MM samples with bone lytic lesion in red, MM samples without lytic lesions in orange.

**Figure 5.**
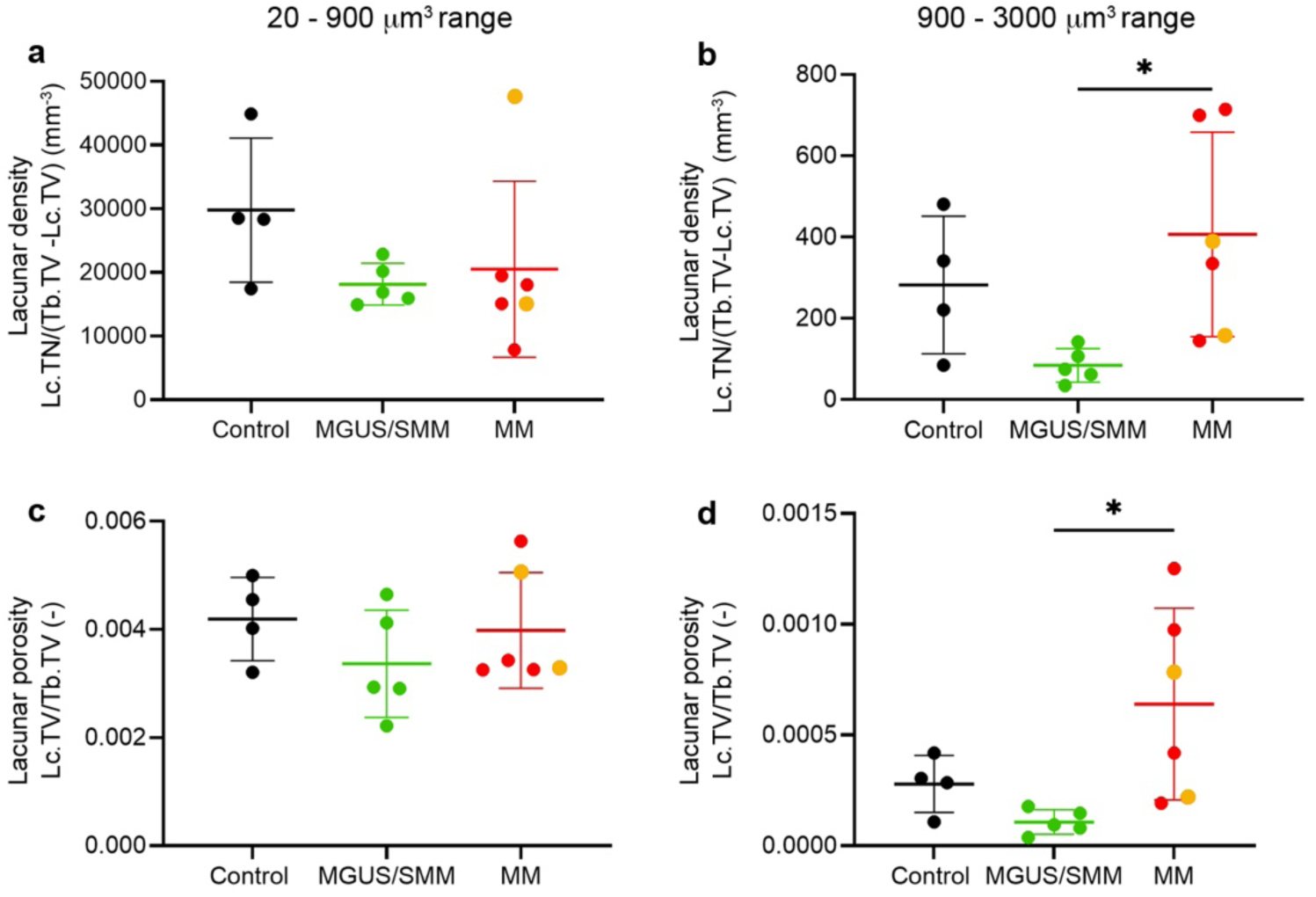
Lacunar density and microporosity analysis for the small lacunae 20-900 µm^3^ range and for the larger lacunae 900-3000 µm^3^ range of osteocytes in control, MGUS, SMM and MM samples. Lacunar density calculated as the total number of osteocyte lacunae (Lc.TN) normalized to the trabecular bone total volume and excluding lacunar total volume (Tb.TV-Lc.TV), for each group control, MGUS/SMM and newly diagnosed MM (samples with bone lytic lesion in red, samples without lytic lesions in orange). Microporosity calculated as lacunar total volume (Lc.TV) divided by trabecular bone total volume (Tb.TV) for each group. Error bars indicate mean ± SD. One way ANOVA with Tukey’s multiple comparison test *p<0.05.

**Figure 6.**
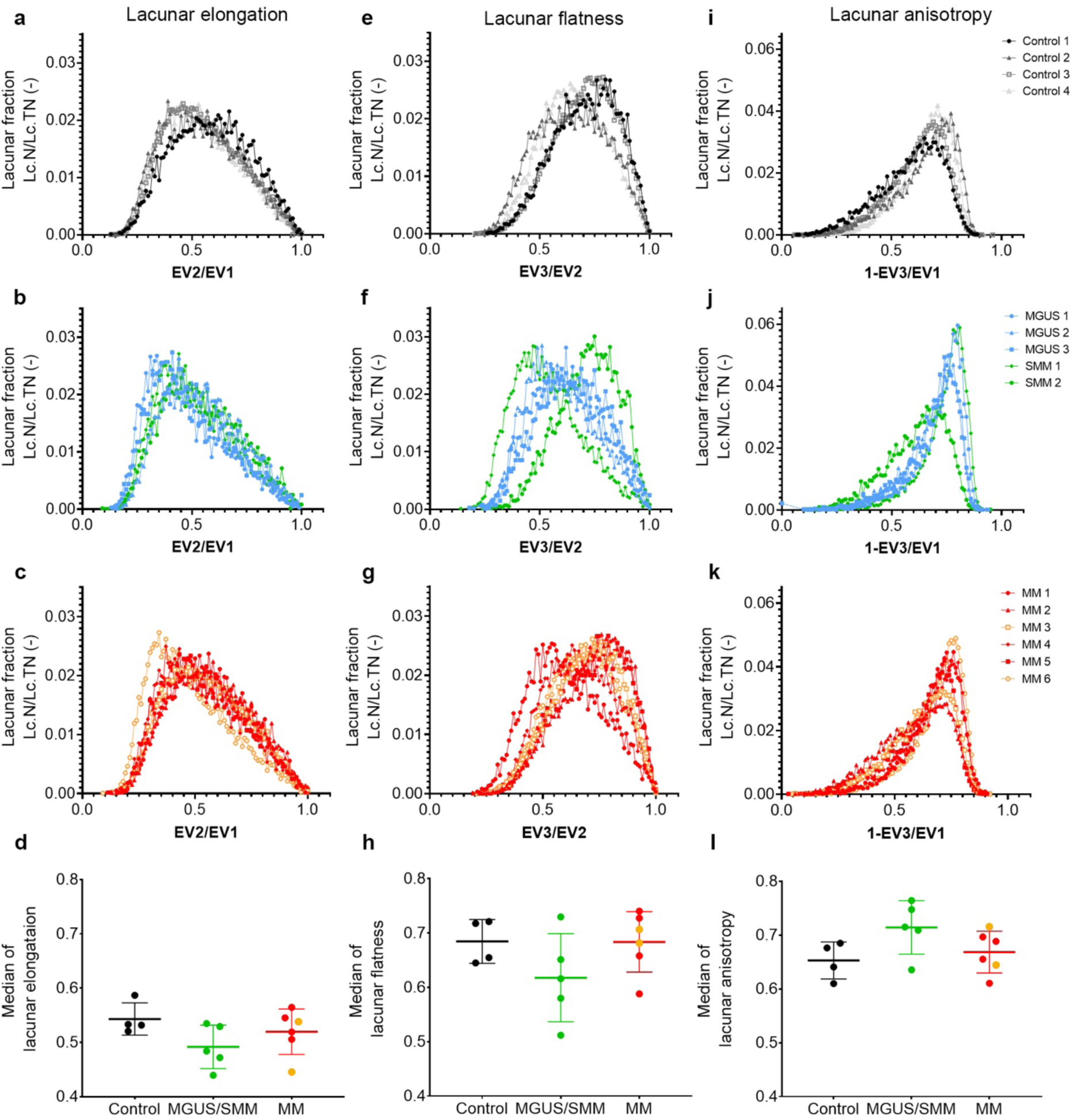
Lacunar morphological analysis for the small lacunae 20-900 µm^3^ range of osteocytes in control, MGUS, SMM and MM samples. Lacunae elongation, flatness and anisotropy analysis for (a, e, i) control, (b, f, j) MGUS/SMM and (c, g, k) MM groups. Individual lacunae were assigned with eigenvalues (EV1, EV2 and EV3) to then calculate lacuna elongation as EV2/EV1 (with lower values for elongated and 1 for spherical) as (a-c) lacunar fraction (Lc.N/Lc.TN) and as (d) median of all lacuna elongation values for each sample; lacuna flatness as EV3/EV2 (ranging from 0 for flat to 1 for spherical) as (e-g) lacunar fraction (Lc.N/Lc.TN) and as (h) median of all lacuna flatness values for each sample; and lacuna anisotropy as 1-EV3/EV1 (with smaller values for isotropic/spherical and 1 for highest anisotropy) as (i-k) lacunar fraction (Lc.N/Lc.TN) and as (l) median of all lacuna anisotropy values for each sample. Sample from the Jamshidi biopsy in black, all other control samples in grey. MGUS in blue, SMM in green, MM samples with bone lytic lesion in red, MM samples without lytic lesions in orange. Error bars indicate mean ± SD.

### 3.4. Analysis of osteocyte lacunar volume distribution and microporosity

First, we characterized the osteocyte lacunar volume distribution, first as a frequency distribution (Fig. 3) and then as a cumulative distribution (Fig. 4). To plot the lacunar volume frequency distribution, all lacunae within one sample were stratified according to volume intervals of 20 μm^3^. Two separate lacunar volume ranges were investigated: small lacunae 20-900 µm^3^ (Fig. 3a-c) (osteocyte lacunar microporosity characteristic of healthy bone ^28,31^), and larger lacunae 900-3000 µm^3^ (Fig. 3d-f) (microporosity potentially associated to bone disease). Control samples (Fig. 3a, d) were compared with MGUS, SMM (Fig. 3b, e), and newly diagnosed MM (Fig. 3c, f). In the small lacunae range 20-900 µm^3^, we observed a distribution within a comparable range (y-axis from 0-4000 µm^3^) in the different groups, with the exception of sample #MM 3, which had a much higher number of osteocyte lacunae compared to the other samples within the MM group (Fig. 3c, note the extended y-axis). In contrast, in the larger lacunae range 900-3000 µm^3^, we observe a clearly higher number of lacunae for the MM group (Fig. 3f) compared with MGUS (Fig. 3d) and SMM (Fig. 3e) groups.

When comparing both ranges, we observed a larger number of small osteocyte lacunae (Fig. 3a-c) compared to larger lacunae (Fig. 3d-f). This motivated a more detailed lacunar volume cumulative distribution analysis (Fig. 4), where the lacunar number (Lc.N) was plotted as a cumulative curve. The 50% cutoff indicates that half of all osteocyte lacunae within one sample are below that specific volume interval (x-axis). In the control group (Fig. 4a), the 50% cutoff is heterogeneous: samples #control 1, #control 3 and #control 4 have 50% of the lacunae with volumes around 140-170 μm^3^, while sample #control 2 has the highest volume with 270 μm^3^. In the MGUS/SMM group (Fig. 4b), the 50% cutoff ranges from 170-210 μm^3^, except sample #SMM 2 with the smallest cutoff at 140 μm^3^. In the MM group (Fig. 4c), the 50% cutoff varies from 150-220 μm^3^, with clearly lower values for the biopsies of patients without bone osteolytic lesions (#MM 3 and #MM 6). In conclusion, we observe a trend for increasing 50% cutoff for more advanced disease, indicating that the sum volume of the smallest 50% lacunae account for a larger volume as the disease progresses (Fig. 4c). In other words, we observe a shift towards larger lacunae with disease progression.

Next, we calculated the lacunar density as the total number of osteocyte lacunae (Lc.TN) normalized to the trabecular bone total volume and excluding lacunar total volume (Tb.TV- Lc.TV), in other words, the sum of the osteocyte lacunar volume frequency distribution (Fig. 3). This was done for both ranges separately, small lacunae 20-900 µm^3^ (Fig. 5a) and larger lacunae 900-3000 µm^3^ (Fig. 5b). In the small lacunae range, the mean for the control samples was 29809 +/- 11314 lacunae/mm^3^, for MGUS/SMM 18165 +/- 3278 lacunae/mm^3^ and for MM 20502 +/- 13827 lacunae/mm^3^. No significant differences were observed in the small lacunae range, although there was a clear trend for lower lacunar density for diseased samples (MGUS/SMM and MM) compared to the controls. However, in the larger lacunae range 900- 3000 µm^3^ (Fig. 5b), significantly higher lacunar density was found in the newly diagnosed MM group compared to the MGUS/SMM group.

In order to evaluate the microporosity in relation to the total bone volume, we divided the lacunar total volume (Lc.TV) by trabecular bone total volume (Tb.TV) for each group. This was done for both ranges separately, small lacunae 20-900 µm^3^ (Fig. 5c) and larger lacunae 900-3000 µm^3^ (Fig. 5d). No significant differences were observed in the small lacunae range. However, in the larger lacunae range, significantly higher microporosity was found in the newly diagnosed MM group compared to the MGUS/SMM group. This is consistent with the observations of lacunar volume frequency distribution (Fig. 3f) and lacunar density (Fig. 5b).

### 3.5 Analysis of osteocyte lacunar morphology

Next, we aimed to assess whether disease progression could be associated with lacunar morphological changes. To do so, we performed a lacunar morphological analysis in the small lacunae 20-900 µm^3^ range in all groups (control, MGUS/SMM and newly diagnosed MM). Based on previous analogous analysis ^34^, we investigated the distribution of lacunar flatness, elongation and anisotropy. Towards that end, the principal eigenvalues for each osteocyte lacunae were obtained: EV1, EV2 and EV3. The morphological parameters were calculated as described in Table 6. The lacuna elongation (EV2/EV1, ratio of medium to largest eigenvalue) is shown in Fig. 6a-d; lacuna flatness (EV3/EV2, ratio of smallest to medium eigenvalue) in Fig. 6e-h; and lacuna anisotropy (1-EV3/EV1, one minus the ratio of smallest to largest eigenvalue) in Fig. 6i-l. The fraction of lacunar number (Lc.N) normalized against lacunae total number (Lc.TN) in each sample is shown for each group: control (Fig. 6a, e, i), MGUS/SMM (Fig. 6b, f, j) and MM (Fig. 6c, g, k). The median of all lacuna elongation values for each sample are shown in Fig. 6d; similarly, for flatness in Fig. 6h and anisotropy in Fig. 6l.

Lacuna flatness in MGUS/SMM and newly diagnosed MM groups exhibit a broader spread of the distribution compared to the control group, with a shift towards lower values (flatter shapes). Regarding lacuna elongation and anisotropy, the spread of the distribution for the three groups is comparable. A trend for lower values for lacuna elongation (more elongated) and higher values for lacuna anisotropy (more anisotropic) is observed in the MGUS/SMM group compared to the control group, as shown by the curves and the by the median values. However, the differences of the median values are not significant.

## 4. Discussion

Changes in the volume and shape of osteocytes were identified in previous studies of bone metastasis from breast and prostate cancer and were indicators of osteolysis ^13–16^. However, high resolution microCT characterization of osteocyte lacunae in human bone biopsies of MGUS, SMM and newly diagnosed MM has not yet been performed, nor have changes in morphology of osteocytes been linked to patterns of osteolysis and disease progression. In this context, quantitative analyses of microCT demonstrate altered lacunae volume and shape distribution between MGUS, SMM and newly diagnosed MM. Specifically, we found a significantly higher number of larger lacunae in the 900-3000 µm^3^ range, in the MM group compared to the MGUS/SMM group, as reflected in higher lacunar volume frequency distribution (Fig. 3f) and significantly higher lacunar density (Fig. 5b). Consistent with the observation of an increase in lacunar volume frequency distribution, we also found significantly higher microporosity for the larger lacunae range in the MM group compared to the MGUS/SMM group (Fig. 5d).

In the small lacunae 20-900 µm^3^ range, the lacunar density for control samples with 29809 +/- 11314 lacunae/mm^3^ (Fig. 5a) is in the range previously reported in the literature for 3D analysis of osteocyte morphometry in healthy transiliac bone biopsies ^28^, which confirms the validity of our analysis. Interestingly, there was a trend for lower lacunar density for diseased samples (MGUS/SMM and MM) compared to the controls, though the differences were not significant (Fig. 5a). This could be indicative of micropetrosis, i.e. osteocyte lacunae mineralization following osteocyte apoptosis ^37,38^. Since a lower lacunar density can distinguish healthy, osteoporotic and bisphosphonate-treated osteoporotic patients, it is currently being considered as a novel bone microstructural marker of impaired bone quality ^37^. In addition, we observed a trend for increasing values of the 50% cutoff in the lacunar volume cumulative distribution for more advanced disease, showing that the sum volume of the smallest 50% lacunae account for a larger volume as the disease progresses (Fig. 4). In terms of shape distribution, the MGUS/SMM group showed a trend for lower lacuna flatness (flatter), lower lacuna elongation (more elongated), and higher lacuna anisotropy (more anisotropic) compared to the control group (Fig. 6). Altogether, the findings of a trend for lower lacunar density and a shift towards larger lacunae (increasing 50% cutoff of the lacunar volume cumulative distribution) for diseased samples diseased samples (MGUS/SMM and MM) compared to control, in the small lacunae 20-900 μm^3^ range; together with significantly higher lacunar density and microporosity for the newly diagnosed MM group, in the larger lacunae 900-3000 μm^3^ range, suggest that the osteocytes in MM bone disease and the precursor conditions MGUS/SMM are undergoing changes in their lacunae morphology during disease progression.

In relation to previous work, currently this is the first study evaluating human bone biopsies at the microstructural level, and so we can only refer to similar work in preclinical setting. Indeed, our established mouse model of MM bone disease was assessed for ultrastructural characterization of the mineralized and nonmineralized ECM, particularly of the osteocyte LCN in early stages of the disease ^27,39–41^. Skeletally mature mice were injected with MM cells and the osteocyte LCN architecture was analyzed using advanced imaging techniques, such as high resolution microCT, synchrotron phase contrast-enhanced microCT and confocal laser scanning microscopy (CLSM) ^27^. Similar as in human samples, we identified alterations in the osteocyte morphology in the early MM lesions and showed larger, irregular-shaped osteocyte lacunae within a disorganized canaliculi network, compared to flat osteocyte lacunae organized in a lamellar structure in healthy bone ^27^. MM affected regions revealed that the lacunae volume was greater and the density was sparser compared to control bones ^27^. These data are in concordance with the human MM samples in which a higher number of larger lacunae and a lower lacunar density were detected in the MM group compared to the precursor conditions and control samples. Overall, the MM preclinical model served to predict the changes in the osteocyte lacunae, which in human bone biopsies occurred as a result of progression from MGUS to MM, and thus aligning the preclinical observations with the human findings.

In a recent study in immunocompetent mice, breast and prostate cancer cells were intratibially injected and the changes in the osteocyte LCN induced by tumor cells were analyzed by histology, microCT imaging and CLSM ^20^. Interestingly, large lacunae and vascular canals were not observed in the proximity of osteolytic lesions, whereas these changes occurred in osteosclerotic lesions adjacent to prostate cancer cells ^20^, indicating that changes of osteocyte lacunae characteristics are tumor type-specific. In our study of human biopsy samples with newly diagnosed MM and the precursor conditions, we observed a significantly higher lacunar density in the MM group compared to the MGUS/SMM groups. These observed changes may be related to the presence of osteocytic osteolysis ^42^, a process that involves osteocytes mobilizing calcium and phosphate, using carbonic anhydrase and vacuolar ATPase to demineralize the bone matrix, and digest protein using matrix metalloproteases and tartrate-resistant acid phosphatase (TRAP) ^20,43–45^. In the above-mentioned mouse model ^20^, TRAP-positive osteocytes in combination with large lacunae were detected in osteosclerotic regions of breast and prostate infiltrated bones, indicating that this mechanism could contribute to the observed increase in lacunae volume ^20,43–45^. Whether osteocytic osteolysis plays a role in the progression from healthy to MGUS/SMM to MM bone is not known and warrants further studies. Moreover, since the osteocyte remodeling function affects the lacunae morphology and the LCN architecture, it is likely that changes in the latter are indicators for impaired bone remodeling activity and mechanoresponsiveness.

In order to understand the implications of altered lacunae volume and shape distribution in MM, future studies should consider several factors. First, there is a need for more bone biopsies in i) each of the groups and ii) with sufficient quality to obtain a larger sample size and validate the findings of this study, iii) at other sites of the skeleton where imaging, for example via PET-CT, has shown infiltration of the bone, and iv) from males and females at the ratio of the occurrence of the disease. Second, it is important that the bone biopsies are obtained from comparable anatomical sites to minimize the natural osteocyte volume and shape variation within healthy bone ^8,28,46^. Third, although previous studies have used a comparable voxel size of 1.2 μm for lacunar imaging in human bone biopsies ^11,28^, to further investigate the relationship between osteocyte lacunae and MM, researchers could increase the volume of interest scanned at high resolution (1 µm voxel size or lower) using synchrotron phase-contrast microCT. Fourth, there is a lack of detailed information whether the biopsies were extracted from bone regions affected or unaffected by the disease. Therefore, it remains uncertain whether the evaluated osteocytes accurately reflect the disease stage. For ethical and practical considerations, the bone samples used in the present study were extracted through a standardized approach from the posterior superior iliac crest. Finally, the use of mouse models of MM may provide a valuable tool for investigating the mechanisms underlying the observed alterations in osteocyte lacunae morphology and other aspects of bone ultrastructure, such as LCN architecture, bone composition, and micro- and nanoscale organization of the organic and inorganic matrix. Furthermore, these models can be used to study how alterations of the osteocyte lacunar morphology affect the response of osteocytes to local tissue strains, their remodeling capacity of mineralized bone matrix and the associated resistance to fractures.

Taken together, these findings suggest that altered lacunae volume and shape distribution are indicative for osteolysis and disease progression from MGUS, SMM to newly diagnosed MM. Future studies are needed to explore whether such changes cause lower local tissue strains and an impaired bone mechanoresponse in MM patients with osteolytic bone lesions and an associated high fracture risk.

## 6. Data availability

The raw/processed data required to reproduce these findings are available in the publicly accessible Edmond repository of the Max Planck Society (https://doi.org/10.17617/3.5BPJUP).

## 7. Author contributions

A Cipitria, I. Moreno-Jiménez and F. Jundt conceived the idea. I. Moreno-Jiménez and S. Heinig processed the bone biopsy samples and performed the experiments. D. Maichl collected and prepared the human Jamshidi bone biopsy samples. S. Strifler educated patients about and performed the biopsies, and collected clinical data. E. Leich curated clinical data related to the human bone biopsy samples; S. Blouin provided the control bone autopsy samples. I. Moreno-Jiménez, S. Heinig, U. Heras, F. Jundt and A. Cipitria analyzed the data. P. Fratzl and N. Fratzl-Zelman together with A Cipitria, I. Moreno-Jiménez and F. Jundt evaluated the methods and results. A Cipitria, I. Moreno-Jiménez and F. Jundt drafted the manuscript. All authors discussed the results and contributed to the final manuscript.

## 9. Acknowledgements

The authors thank Daniel Werner, Jeannette Steffen, Daniela S. Garske and Susann Weichold at the Max Planck Institute of Colloids and Interfaces in Potsdam for their technical assistance.

## References

1. Bonewald, L. F. The amazing osteocyte. Journal of Bone and Mineral Research 26, 229–238 (2011).

2. Hemmatian, H., Bakker, A. D., Klein-Nulend, J. & van Lenthe, G. H. Aging, osteocytes, and mechanotransduction. Curr Osteoporos Rep 15, 401–411 (2017).

3. Gerbaix, M., Gnyubkin, V., Farlay, D., Olivier, C., Ammann, P., Courbon, G., Laroche, N., Genthial, R., Follet, H., Peyrin, F., Shenkman, B., Gauquelin-Koch, G. & Vico, L. One-month spaceflight compromises the bone microstructure, tissue-level mechanical properties, osteocyte survival and lacunae volume in mature mice skeletons. Sci Rep 7, 2659 (2017).

4. Lane, N. E., Yao, W., Balooch, M., Nalla, R. K., Balooch, G., Habelitz, S., Kinney, J. H. & Bonewald, L. F. Glucocorticoid-treated mice have localized changes in trabecular bone material properties and osteocyte lacunar size that are not observed in placebo-treated or estrogen-deficient mice. Journal of Bone and Mineral Research 21, 466–476 (2005).

5. Tiede-Lewis, L. M. & Dallas, S. L. Changes in the osteocyte lacunocanalicular network with aging. Bone 122, 101–113 (2019).

6. Tommasini, S. M., Trinward, A., Acerbo, A. S., De Carlo, F., Miller, L. M. & Judex, S. Changes in intracortical microporosities induced by pharmaceutical treatment of osteoporosis as detected by high resolution micro-CT. Bone 50, 596–604 (2012).

7. Mullender, M. G., van der Meer, D. D., Huiskes, R. & Lips, P. Osteocyte density changes in aging and osteoporosis. Bone 18, 109–113 (1996).

8. Blouin, S., Misof, B. M., Mähr, M., Fratzl-Zelman, N., Roschger, P., Lueger, S., Messmer, P., Keplinger, P., Rauch, F., Glorieux, F. H., Berzlanovich, A., Gruber, G. M., Brugger, P. C., Shane, E., Recker, R. R., Zwerina, J. & Hartmann, M. A. Osteocyte lacunae in transiliac bone biopsy samples across life span. Acta Biomater 157, 275–287 (2023).

9. Milovanovic, P., Zimmermann, E. A., Riedel, C., Scheidt, A. vom, Herzog, L., Krause, M., Djonic, D., Djuric, M., Püschel, K., Amling, M., Ritchie, R. O. & Busse, B. Multi- level characterization of human femoral cortices and their underlying osteocyte network reveal trends in quality of young, aged, osteoporotic and antiresorptive-treated bone. Biomaterials 45, 46–55 (2015).

10. Hemmatian, H., Bakker, A. D., Klein-Nulend, J. & van Lenthe, G. H. Alterations in osteocyte lacunar morphology affect local bone tissue strains. J Mech Behav Biomed Mater 123, 104730 (2021).

11. Yang, K. G., Goff, E., Cheng, K. lo, Kuhn, G. A., Wang, Y., Cheng, J. C. yiu, Qiu, Y., Müller, R. & Lee, W. Y. wai. Abnormal morphological features of osteocyte lacunae in adolescent idiopathic scoliosis: A large-scale assessment by ultra-high-resolution micro-computed tomography. Bone 166, 116594 (2023).

12. Atkinson, E. G. & Delgado-Calle, J. The emerging role of osteocytes in cancer in bone. JBMR Plus 3, e10186 (2019).

13. Adhikari, M. & Delgado-Calle, J. Role of osteocytes in cancer progression in the bone and the associated skeletal disease. Curr Osteoporos Rep 19, 247–255 (2021).

14. Hofbauer, L. C., Bozec, A., Rauner, M., Jakob, F., Perner, S. & Pantel, K. Novel approaches to target the microenvironment of bone metastasis. Nat Rev Clin Oncol 18, 488–505 (2021).

15. Mastro, A. M., Gay, C. V., Welch, D. R., Donahue, H. J., Jewell, J., Mercer, R., DiGirolamo, D., Chislock, E. M. & Guttridge, K. Breast cancer cells induce osteoblast apoptosis: A possible contributor to bone degradation. J Cell Biochem 91, 265–276 (2004).

16. Hemmatian, H., Conrad, S., Furesi, G., Mletzko, K., Krug, J., Faila, A. V., Kuhlmann, J. D., Rauner, M., Busse, B. & Jähn-Rickert, K. Reorganization of the osteocyte lacuno-canalicular network characteristics in tumor sites of an immunocompetent murine model of osteotropic cancers. Bone 152, 116074 (2021).

17. Roodman, G. Pathogenesis of myeloma bone disease. Blood Cells Mol Dis 32, 290–292 (2004).

18. Berenson, J., Rajdev, L. & Broder, M. Bone complications in multiple myeloma. Cancer Biol Ther 5, 1082–1085 (2006).

19. Terpos, E., Ntanasis-Stathopoulos, I. & Dimopoulos, M. A. Myeloma bone disease: from biology findings to treatment approaches. Blood 133, 1534–1539 (2019).

20. Delgado-Calle, J., Anderson, J., Cregor, M. D., Hiasa, M., Chirgwin, J. M., Carlesso, N., Yoneda, T., Mohammad, K. S., Plotkin, L. I., Roodman, G. D. & Bellido, T. Bidirectional notch signaling and osteocyte-derived factors in the bone marrow microenvironment promote tumor cell proliferation and bone destruction in multiple myeloma. Cancer Res 76, 1089–1100 (2016).

21. Huston, A. & Roodman, G. D. Role of the microenvironment in multiple myeloma bone disease. Future Oncology 2, 371–378 (2006).

22. Rajkumar, S. V., Dimopoulos, M. A., Palumbo, A., Blade, J., Merlini, G., Mateos, M.- V., Kumar, S., Hillengass, J., Kastritis, E., Richardson, P., Landgren, O., Paiva, B., Dispenzieri, A., Weiss, B., LeLeu, X., Zweegman, S., Lonial, S., Rosinol, L., Zamagni, E., Jagannath, S., Sezer, O., Kristinsson, S. Y., Caers, J., Usmani, S. Z., Lahuerta, J. J., Johnsen, H. E., Beksac, M., Cavo, M., Goldschmidt, H., Terpos, E., Kyle, R. A., Anderson, K. C., Durie, B. G. M. & Miguel, J. F. S. International myeloma working group updated criteria for the diagnosis of multiple myeloma. Lancet Oncol 15, e538–e548 (2014).

23. Borggrefe, J., Giravent, S., Campbell, G., Thomsen, F., Chang, D., Franke, M., Günther, A., Heller, M. & Wulff, A. Association of osteolytic lesions, bone mineral loss and trabecular sclerosis with prevalent vertebral fractures in patients with multiple myeloma. Eur J Radiol 84, 2269–2274 (2015).

24. Lee, E. M. & Kim, B. Clinical significance of trabecular bone score for prediction of pathologic fracture risk in patients with multiple myeloma. Osteoporos Sarcopenia 4, 73–76 (2018).

25. Takasu, M., Tani, C., Ishikawa, M., Date, S., Horiguchi, J., Kiguchi, M., Tamura, A., Sakai, A., Asaoku, H., Nango, N. & Awai, K. Multiple Myeloma: Microstructural Analysis of Lumbar Trabecular Bones in Patients without Visible Bone Lesions— Preliminary Results. Radiology 260, 472–479 (2011).

26. Michels, M., Morais-Faria, K., Rivera, C., Brandão, T. B., Santos-Silva, A. R. & Oliveira, M. L. Structural complexity of the craniofacial trabecular bone in multiple myeloma assessed by fractal analysis. Imaging Sci Dent 52, 33–41 (2022).

27. Ziouti, F., Soares, A. P., Moreno-Jiménez, I., Rack, A., Bogen, B., Cipitria, A., Zaslansky, P. & Jundt, F. An early myeloma bone disease model in skeletally mature mice as a platform for biomaterial characterization of the extracellular matrix. J Oncol 2020, 1–12 (2020).

28. Goff, E., Cohen, A., Shane, E., Recker, R. R., Kuhn, G. & Müller, R. Large-scale osteocyte lacunar morphological analysis of transiliac bone in normal and osteoporotic premenopausal women. Bone 160, 116424 (2022).

29. Carter, Y., Thomas, C. D. L., Clement, J. G. & Cooper, D. M. L. Femoral osteocyte lacunar density, volume and morphology in women across the lifespan. J Struct Biol 183, 519–526 (2013).

30. Buenzli, P. R. & Sims, N. A. Quantifying the osteocyte network in the human skeleton. Bone 75, 144–150 (2015).

31. McCreadie, B. R., Hollister, S. J., Schaffler, M. B. & Goldstein, S. A. Osteocyte lacuna size and shape in women with and without osteoporotic fracture. J Biomech 37, 563–572 (2004).

32. Gescheidtova, E., Marcon, P., Bartusek, K., Gescheidtová, E., Marcon, P. & Bartusek, K. Visualization of plant fibres via diffusion tensor imaging. in PIERS Proceedings, Suzhou, China (2011). at <https://www.researchgate.net/profile/Karel-Bartusek/publication/267556847_Visualization_of_Plant_Fibres_via_Diffusion_Tensor_Imaging/links/5583ec1a08ae89172b861a80/Visualization-of-Plant-Fibres-via-Diffusion-Tensor-Imaging.pdf>

33. Mader, K. S., Schneider, P., Müller, R. & Stampanoni, M. A quantitative framework for the 3D characterization of the osteocyte lacunar system. Bone 57, 142–154 (2013).

34. Chaumel, J., Schotte, M., Bizzarro, J. J., Zaslansky, P., Fratzl, P., Baum, D. & Dean, M. N. Co-aligned chondrocytes: Zonal morphological variation and structured arrangement of cell lacunae in tessellated cartilage. Bone 134, 115264 (2020).

35. Bernstein, Z. S., Kim, E. B. & Raje, N. Bone Disease in Multiple Myeloma: Biologic and Clinical Implications. Cells 2022, Vol. 11, Page 2308 11, 2308 (2022).

36. Palumbo, A., Avet-Loiseau, H., Oliva, S., Lokhorst, H. M., Goldschmidt, H., Rosinol, L., Richardson, P., Caltagirone, S., Lahuerta, J. J., Facon, T., Bringhen, S., Gay, F., Attal, M., Passera, R., Spencer, A., Offidani, M., Kumar, S., Musto, P., Lonial, S., Petrucci, M. T., Orlowski, R. Z., Zamagni, E., Morgan, G., Dimopoulos, M. A., Durie, B. G. M., Anderson, K. C., Sonneveld, P., San Miguel, J., Cavo, M., Rajkumar, S. V. & Moreau, P. Revised International Staging System for Multiple Myeloma: A Report From International Myeloma Working Group. Journal of Clinical Oncology 33, 2863–2869 (2015).

37. Milovanovic, P. & Busse, B. Phenomenon of osteocyte lacunar mineralization: indicator of former osteocyte death and a novel marker of impaired bone quality? Endocr Connect 9, R70–R80 (2020).

38. Delgado-Calle, J., Bellido, T. & Roodman, G. D. Role of osteocytes in multiple myeloma bone disease. Curr Opin Support Palliat Care 8, 407–413 (2014).

39. Schwarzer, R., Nickel, N., Godau, J., Willie, B. M., Duda, G. N., Schwarzer, R., Cirovic, B., Leutz, A., Manz, R., Bogen, B., Dörken, B. & Jundt, F. Notch pathway inhibition controls myeloma bone disease in the murine MOPC315.BM model. Blood Cancer J 4, e217–e217 (2014).

40. Rummler, M., Ziouti, F., Bouchard, A. L., Brandl, A., Duda, G. N., Bogen, B., Beilhack, A., Lynch, M. E., Jundt, F. & Willie, B. M. Mechanical loading prevents bone destruction and exerts anti-tumor effects in the MOPC315.BM.Luc model of myeloma bone disease. Acta Biomater 119, 247–258 (2021).

41. Ziouti, F., Rummler, M., Steyn, B., Thiele, T., Seliger, A., Duda, G. N., Bogen, B., Willie, B. M. & Jundt, F. Prevention of bone destruction by mechanical loading is not enhanced by the Bruton’s tyrosine kinase inhibitor CC-292 in myeloma bone disease. Int J Mol Sci 22, 3840 (2021).

42. Tsourdi, E., Jähn, K., Rauner, M., Busse, B. & Bonewald, L. F. Physiological and pathological osteocytic osteolysis. J Musculoskelet Neuronal Interact 18, 292–303 (2018).

43. Jähn, K., Kelkar, S., Zhao, H., Xie, Y., Tiede-Lewis, L. M., Dusevich, V., Dallas, S. L. & Bonewald, L. F. Osteocytes acidify their microenvironment in response to PTHrP in vitro and in lactating mice in vivo. Journal of Bone and Mineral Research 32, 1761–1772 (2017).

44. Kogawa, M., Wijenayaka, A. R., Ormsby, R. T., Thomas, G. P., Anderson, P. H., Bonewald, L. F., Findlay, D. M. & Atkins, G. J. Sclerostin regulates release of bone mineral by osteocytes by induction of carbonic anhydrase 2. Journal of Bone and Mineral Research 28, 2436–2448 (2013).

45. Tang, S. Y., Herber, R.-P., Ho, S. P. & Alliston, T. Matrix metalloproteinase-13 is required for osteocytic perilacunar remodeling and maintains bone fracture resistance. Journal of Bone and Mineral Research 27, 1936–1950 (2012).

46. Schemenz, V., Gjardy, A., Chamasemani, F. F., Roschger, A., Roschger, P., Zaslansky, P., Helfen, L., Burghammer, M., Fratzl, P., Weinkamer, R., Brunner, R., Willie, B. M. & Wagermaier, W. Heterogeneity of the osteocyte lacuno-canalicular network architecture and material characteristics across different tissue types in healing bone. J Struct Biol 212, 107616 (2020).

