## Supplementary information for "3D osteocyte lacunar morphometry of human bone biopsies with high resolution microCT: from monoclonal gammopathy to newly diagnosed multiple myeloma"

- Supplementary Figure S1
- Supplementary Figure S2
- Supplementary Figure S3

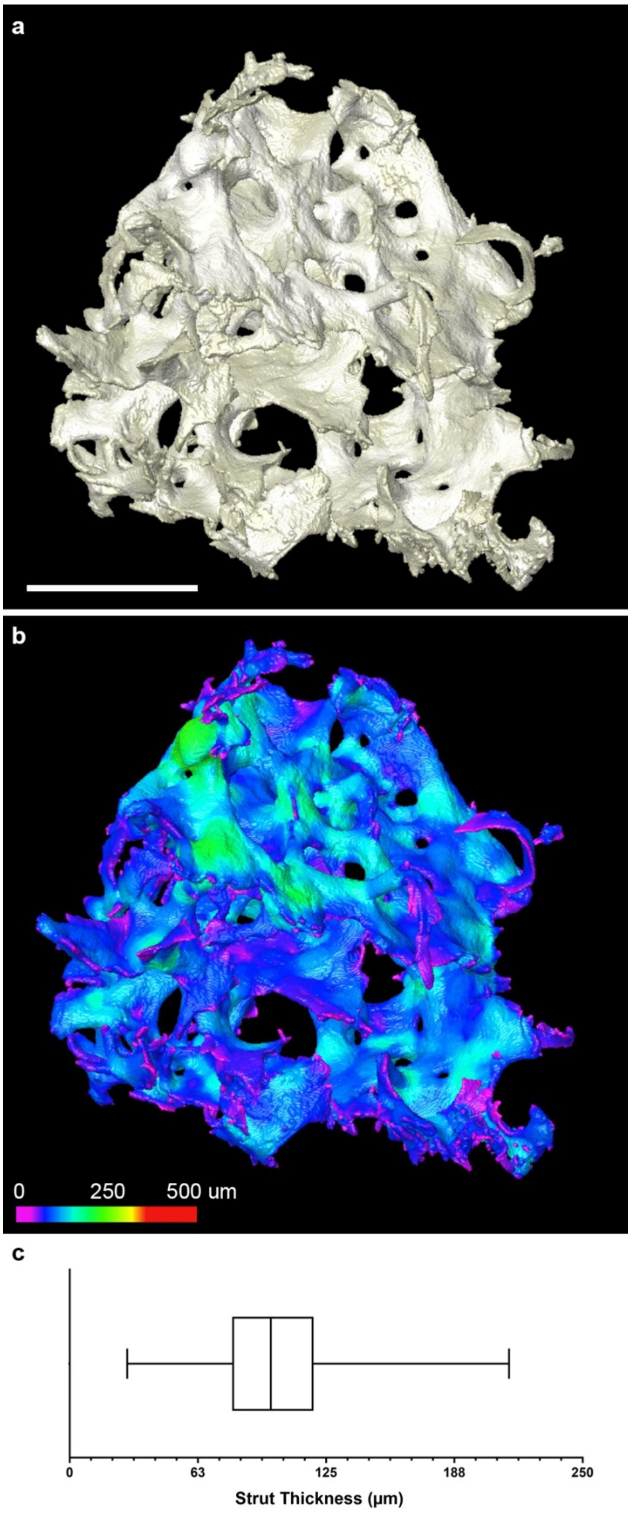


**Supplementary Figure S1. Low resolution microCT overview of a control human bone biopsy.** (a) 3D reconstruction of low resolution (7 µm voxel) microCT scan. (b) Quantitative analysis of trabecular bone strut thickness distribution and color code visualization with Dragonfly software. (c) Distribution of trabecular thickness shown with median, first and third quartile. Scale bars equal to 1 mm.

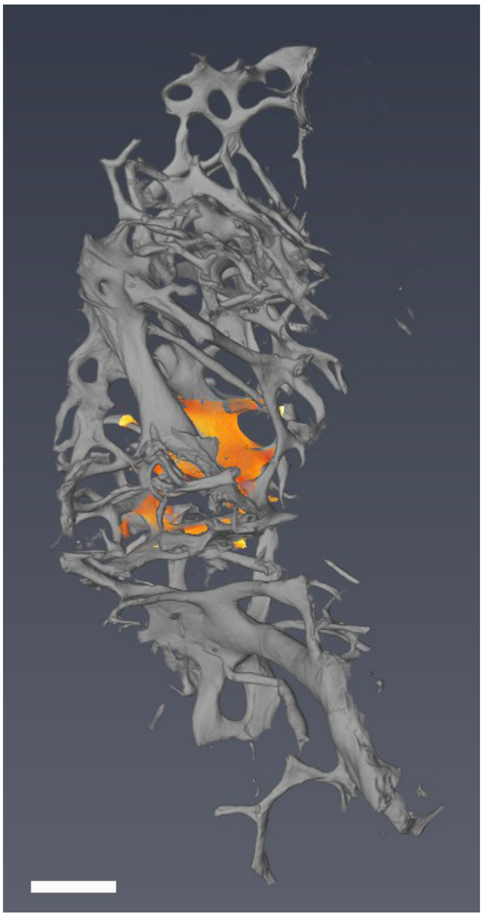

**Supplementary Figure S2.** 3D reconstruction of low resolution (7 μm voxel) microCT scan and representative volume of interest (VOI) for high resolution (1-2 μm voxel) microCT scan (in orange). Scale bar equals to 1 mm.

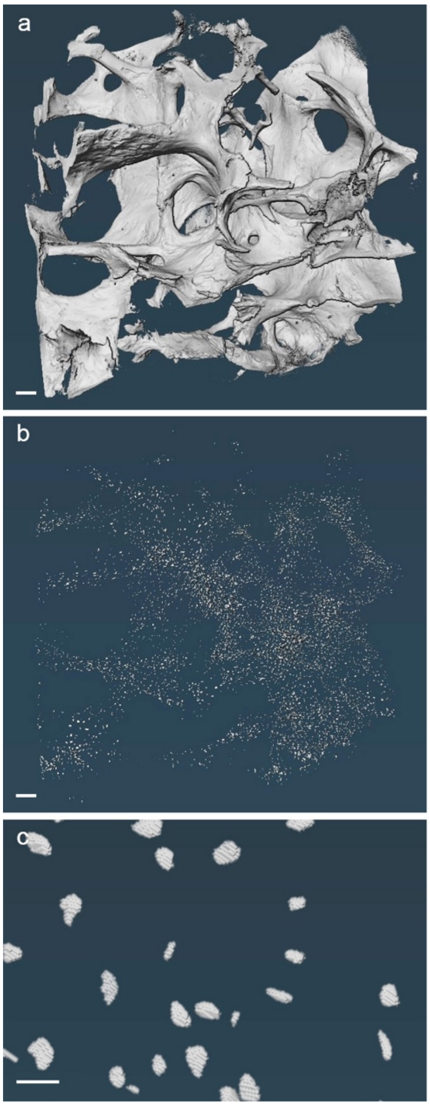


**Supplementary Figure S3. Bone biopsy macroscopic structure and osteocyte lacunar microporosity visualized with high resolution microCT.** (a) 3D reconstruction of local scan (high resolution microCT; voxel size 1-2 µm) and (b-c) the corresponding segmented osteocyte lacunar microporosity of representative samples. Control bone biopsy #1 (Jamshidi). Scale bars equal to 500 µm (a, b) and 20 µm (c).
